# Primate-restricted KRAB zinc finger proteins and target retrotransposons control gene expression in human neurons

**DOI:** 10.1101/856005

**Authors:** Priscilla Turelli, Christopher Playfoot, Dephine Grun, Charlène Raclot, Julien Pontis, Alexandre Coudray, Christian Thorball, Julien Duc, Eugenia Pankevich, Bart Deplancke, Volker Busskamp, Didier Trono

## Abstract

In the first days of embryogenesis, transposable element-embedded regulatory sequences (TEeRS) are silenced by Kruppel-associated box (KRAB)-zinc finger proteins (KZFPs). Many TEeRS are subsequently coopted in transcription networks, but how KZFPs influence this process is largely unknown. We identify ZNF417 and ZNF587 as primate-specific KZFPs repressing HERVK (human endogenous retrovirus K) and SVA (SINE-VNTR-Alu) integrants in human embryonic stem cells (ESC). Expressed in specific regions of the human developing and adult brain, ZNF417/587 keep controlling TEeRS in ESC-derived neurons and brain organoids, secondarily influencing the differentiation and neurotransmission profile of neurons and preventing the induction of neurotoxic retroviral proteins and an interferon-like response. Thus, evolutionarily recent KZFPs and their TE targets partner up to influence human neuronal differentiation and physiology.

**One Sentence Summary:** Young transposable elements and their protein controllers team up to regulate the differentiation and function of human neurons.

## Main Text

Some 4.5 million transposable element (TE)-derived sequences are disseminated across the human genome, many of which integrated within the last few tens of million years (*1*). TEs are typically enriched in transcription factor (TF) binding sites, and correspondingly influence gene expression in a broad range of biological events (*2*–*5*). However, TEeRS are silenced during the earliest phase of embryogenesis by KZFPs, which dock KAP1 (KRAB-associated protein 1, a.k.a. TRIM28) and associated heterochromatin inducers to TE loci (*6*–*8*). The rapid evolutionary selection of KZFP genes was initially interpreted as solely reflecting the host component of an arms race, but recent data suggest that KZFPs team up with TEs to build species-restricted layers of epigenetic regulation (*8, 9*). The present work provides direct support for this model.

We previously determined that a 35bp-long TE sequence encompassing the HERVK14C primer binding site (PBS)-encoding region (coined HERVK-R) confers KAP1-induced repression to a nearby PGK promoter in hESC (*10*). As part of a large-scale screen, we identified ZNF417 and ZNF587 as selectively enriched at loci containing this HERVK sequence (*9*). Depleting these two KZFPs from hESC restored expression of an HERVK-R-containing PGK-GFP lentivector (LV) (**Fig. 1A**), while producing them in murine ESCs silenced this vector, demonstrating the sequence-specific repressor potential of ZNF417 and ZNF587 and the likely absence of mouse orthologue (fig. S1A). Phylogenetic analyses confirmed that *ZNF417* emerged in the human ancestral genome ahead of the New World Monkey split 43 million years ago and that *ZNF587* arose by duplication some 24 million years later (fig. S1, B and C). ZNF417 and ZNF587 display 98% amino acid homology with some differences in their zinc fingerprints, the series of amino acids trios predicted to dictate the sequence specificity of their DNA binding (**Fig. 1B** and fig. S1, B and C). Only rare individuals harbor homozygous loss of function (LoF) mutations in *ZNF417* or *ZNF587* in the gnomAD repertoire (https://gnomad.broadinstitute.org/) (**Fig. 1B**), and the two genes exhibit fairly comparable patterns of expression across tissues according to GTEX (https://gtexportal.org/) and the BrainSpan Atlas of the Developing Human Brain (human.brain-map.org), with higher levels of transcripts in adult pituitary gland, thyroid, ovary, uterus and in pre-natal compared to post-natal brain structures (**Fig. 1C** and fig. S1D).

**Fig. 1:**
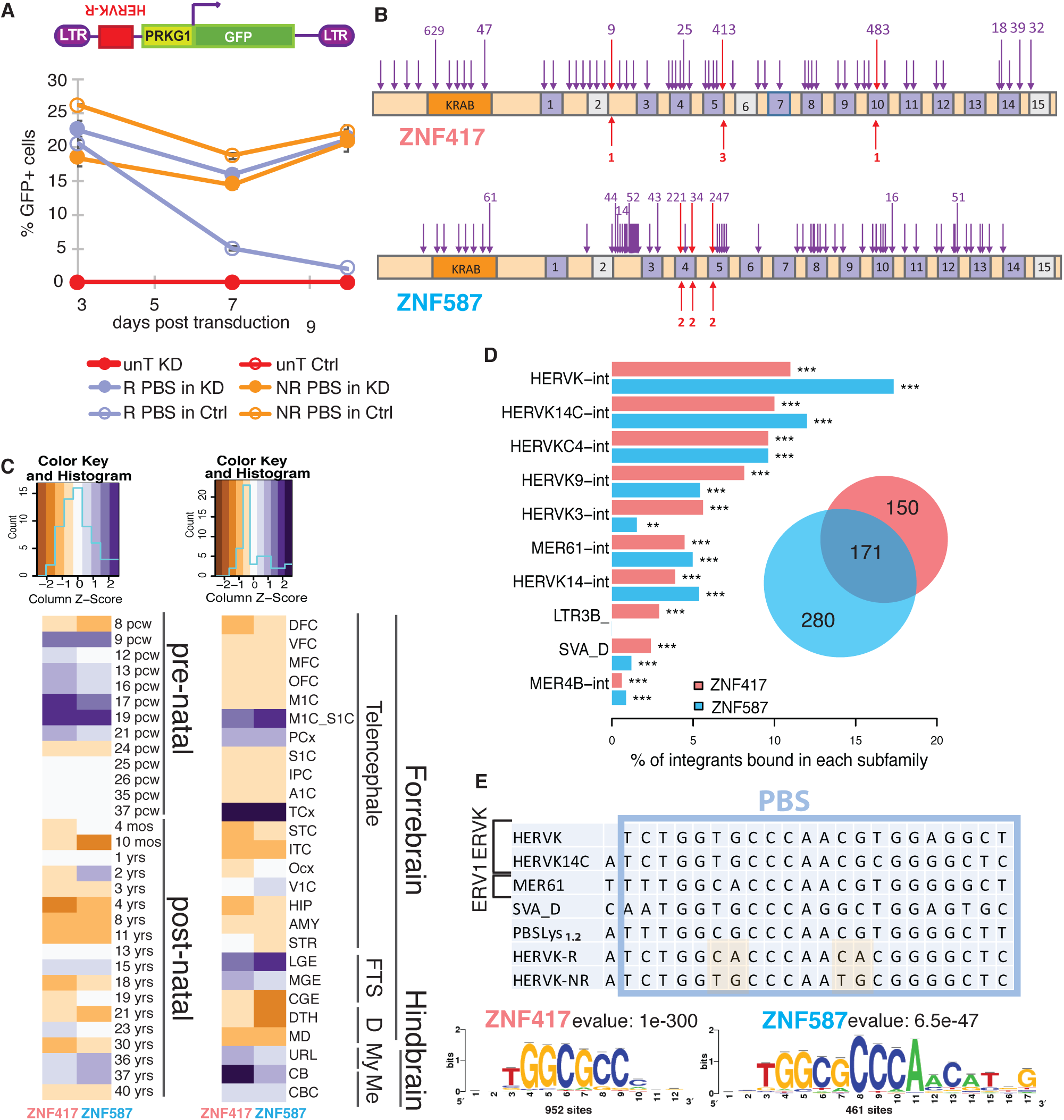
Genomic characterization of ZNF417/587. **(A)** Expression from a PGK-GFP cassette cloned downstream of KAP1-restricted (R) or -non restricted (NR) HERVK PBS sequences in control (Ctrl) or ZNF417/587 KD hESCs (average and s.d. values of duplicates). UnT: untransduced. **(B)** LoF mutations for *ZNF417/587* with numbers of most frequent alleles amongst >140’000 individuals (in red homozygous LoF mutations). Dark and light purple boxes indicate intact and degenerated ZFs. **(C)** ZNF417/587 expression in brain development and substructures according to BrainSpan Atlas of the Developing Human Brain. Pcw, post-conception week. FTS: forebrain fetal transient structures, D: diencephale, My: myelencephale, Me: metencephale, M1C_S1C: primary motor sensory cortex, PCx: parietal cortex, TCx: temporal neocortex, LGE: lateral ganglionic eminence, MGE: medial ganglionic eminence, URL: upper rhombic lip, CB: cerebellum, CBC: cerebellar cortex. **(D)** ZNF417/587-bound TEs in hESCs, with histogram representing percentage of integrants from each TE subfamily (*P* values, hypergeometric test) and Venn diagram total number of ChIP-Seq peaks. **(E)** Top: PBS consensus sequences of bound LTR/ERVs and SVAs, compared with PBS_Lys1.2_ and R- and NR-HERVK14C sequences. Bottom: Predicted binding motifs according to Rsat.

Chromatin immunoprecipitation-sequencing (ChIP-seq) of H1 hESCs overexpressing HA-tagged versions of ZNF417 and ZNF587 identified 321 and 451 peaks, respectively, including 171 in common. About 85% mapped to primate-restricted LTR/ERVK, SVAs and LTR/ERV1 (**Fig. 1D** and fig. S2, A to C), amongst which 12 human-specific LTR/ERVK, and 4 out of 8 HML-2 HERVK previously noted to be polymorphic in the population (*11*) (table S1). KAP1, which binds both KZFPs (fig. S2D), and H3K9me3, the repressive mark instated by the KAP1-associated histone methyltransferase SETDB1, were enriched at the PBS-coding sequence of ZNF417/ZNF587-bound LTR/ERVKs in hESCs (fig. S2E). Most bound ERV sequences correspond to the PBSLys1.2-coding region, and a highly homologous motif is found in SVA-D integrants (**Fig. 1E**). Furthermore, SMILE-seq (*12*) revealed that ZNF417 and ZNF587 had a higher affinity for methylated than unmethylated versions of this sequence (fig. S2F). Correspondingly, these KZFPs inefficiently repressed the HERVK-R-PGK-GFP LV in hESCs depleted for the *de novo* DNA methyltransferases DNMT3A and 3B, although this might also reflect indirect effects (fig. S2G).

The knockdown (KD) of *ZNF417/ZNF587* in hESC resulted in upregulating (fold change>2, FDR<0.05) 857 TEs, most notably members of the LTR/ERV1, LTR/ERVL-MalR, SVA (*P*<0.001, hypergeometric test) and LTR/ERVK (P=0.055) subgroups, many of which harbored a ZNF417/ZNF587 binding motif (**Fig. 2A**, left) and were normally bound by KAP1 (**Fig. 2A** middle) (*P*<0.001, one-sided Fisher’s exact test). Correspondingly, TEs normally bound by these KZFPs lost H3K9me3 and gained H3K4me1 and H3K27ac and were more expressed than their unbound counterparts in knockdown ESC (**Fig. 2B**, top). Expression of 854 genes was also altered (**Fig. 2A**, right, and table S2), a majority upregulated (fold change>2, FDR< 0.05), some (*P*=0.039, one-sided Fisher’s exact test) with a ZNF417/587 peak within 100kb of the transcription start site (TSS). Those within 20kb of a ZNF417/587-bound TE lost H3K9me3 and gained H4K4me1 at the TSS upon KZFP KD, but rarely displayed higher levels of H3K27Ac and transcription, consistent with a poised promoter state (**Fig. 2B**, middle). In contrast, the TSS of genes induced in this setting displayed on average increased levels of H3K4me1 and H3K27ac but no change in H3K9me3 (**Fig. 2B** bottom), probably because many of these genes, including 130 interferon-sensitive genes (ISGs) and the putative targets of 31 upregulated TFs, were indirectly rather than directly controlled by the KZFPs. Using publicly available data, we found that TE integrants bound by ZNF417/ZNF587 in hESC were induced during embryonic genome activation (EGA), repressed upon naïve-to-primed hESC conversion (**Fig. 3A**), and that genes controlled by these two KZFPs were relatively more expressed during human than macaque EGA (fig. S3A), consistent with our recent proposal that KZFPs tame the transcriptional activity of EGA-promoting TE enhancers (*8*). ZNF417/587-targeted TEs were also more expressed in brain and testis than in other tissues (fig. S3B), and we found 40% of these loci to overlap with regions classified as brain and spinal cord enhancers in the EnhancerAtlas 2.0 (*13*). Accordingly, several genes normally expressed in the brain stood out amongst transcriptional units upregulated in hESC depleted for ZNF417/587. For instance, *AADAT*, the product of which facilitates the synthesis of the neuroprotective kynurenic acid (*14*), and *PRODH*, a gene highly expressed in the brain where it influences GABAergic neurotransmission and previously linked to schizophrenia (*15, 16*), both harbor ZNF417/587-recruiting HERVKs upstream of their promoters and were markedly induced by depletion of these KZFPs (**Fig. 3B**). Correspondingly, levels of *ZNF417/587* transcripts anti-correlate during development and in many regions of the adult brain with those of HERVKs and *PRODH* (**Fig. 3C** and fig. S3, C and D).

**Fig. 2:**
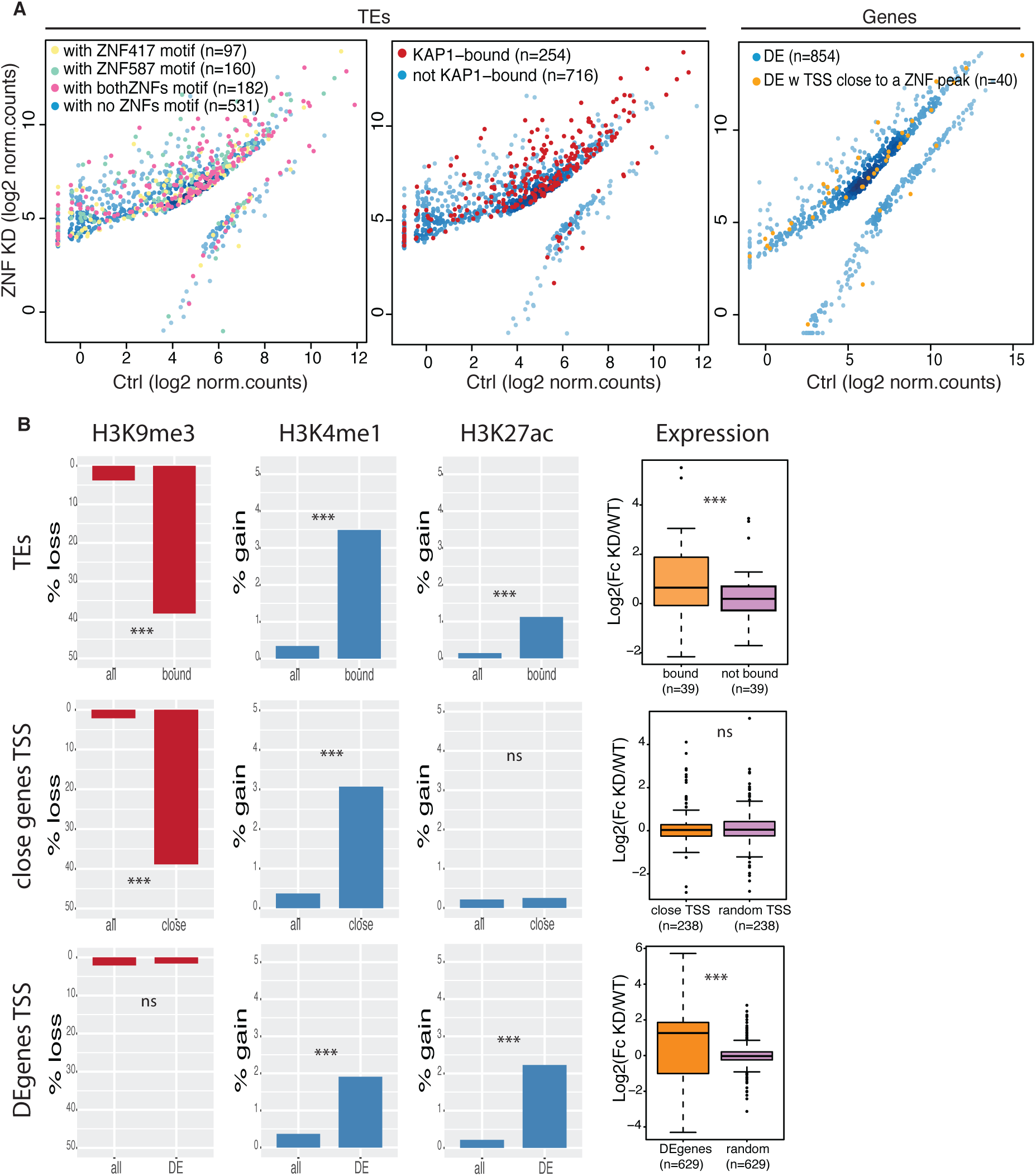
Impact of ZNF417/587 depletion in pluripotent stem cells. **(A)** Dot-plots of RNAseq from ZNF417/587 KD *vs*. control hESCs, outlining differentially expressed (DE) TEs and genes (fold change>2, FDR<0.05). TEs with predicted ZNF417 (yellow), ZNF587 (green) or both (pink) binding motifs (left panel) or bound by KAP1 in hESC (middle panel) or genes with a TSS closer than 100kb from a ZNF binding site (right panel) are highlighted. **(B)** Bar plots depicting loss of H3K9me3 or gain in H3K4me1 or H3K27ac at indicated loci. Upper panel: ZNF417/587-bound *vs*. -unbound TEs; middle panel: TSS of coding genes close to (<20kb) *vs*. distant from a KZFP peak; lower panel: TSS of coding DE genes *vs*. all genes (*P* values, hypergeometric test). Right: fold change in expression in KD *vs*. WT hESC of loci illustrated on left (*P* values, Wilcoxon test).

**Fig. 3:**
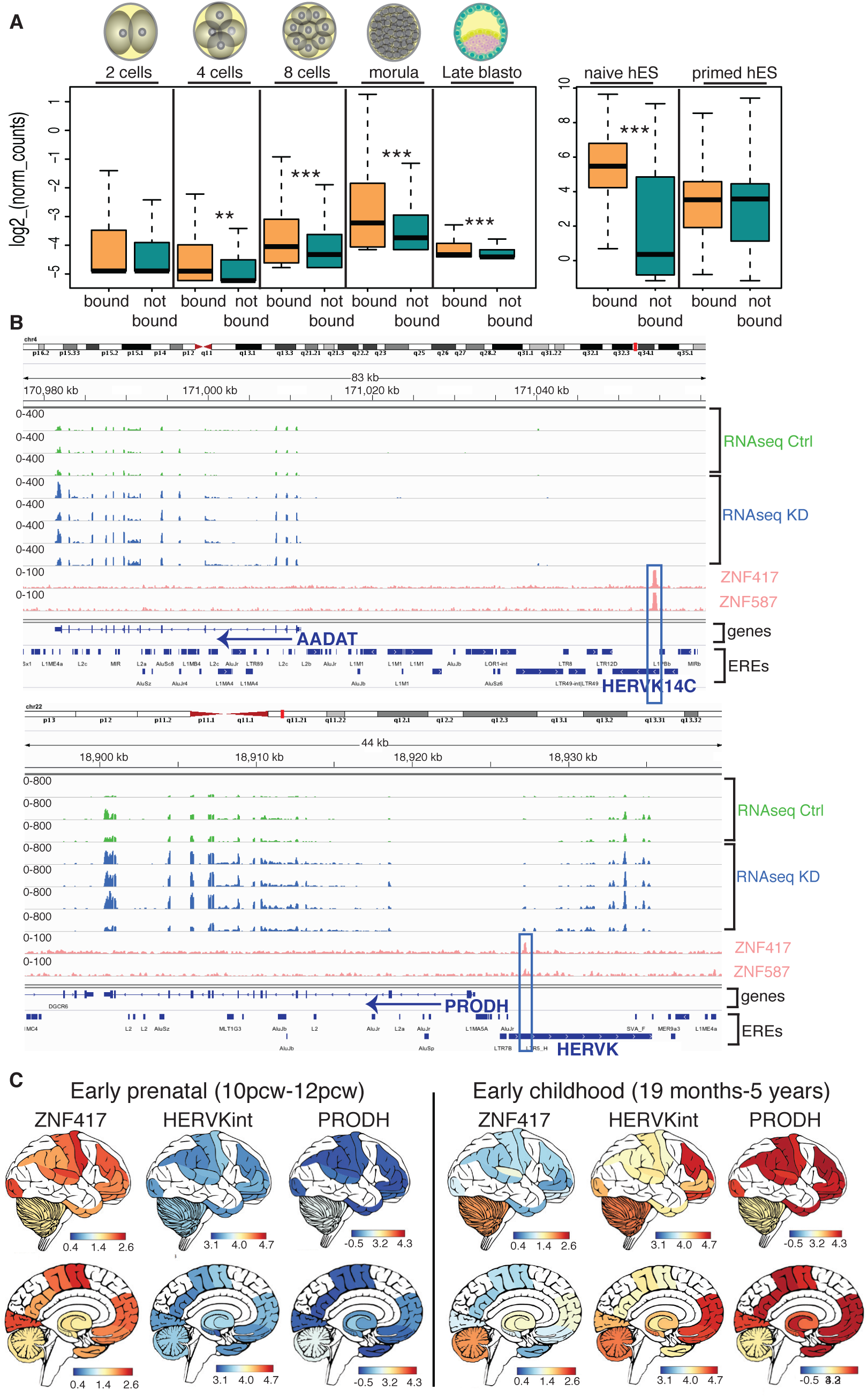
ZNF417/587-mediated repression of TEeRs and neuron-specific genes. **(A)** Expression of TEs bound or not bound at indicated stages of human development using single-cell (left panel) or in naïve versus primed (right panel) hESC RNAseq data. **(B)** IGV screenshots of independent RNAseq replicates from control (Ctrl) and KZFP KD hESCs with boxed ZNF417/587 peaks at HERVK integrants upstream of *AADAT* (upper panel) and *PRODH* (lower panel). **(C)** Spatial representation of *ZNF417, HERVK* and *PRODH* expression in early prenatal and childhood brains, using RNAseqs from the Brain Span Atlas of the Developing Human Brain. Pre-natal brain is depicted as anatomically adult for consistency.

To test functionally whether ZNF417/ZNF587-controlled TEeRS act as neuron-specific enhancers, we first used an *in vitro* neuronal differentiation system where the doxycycline-inducible expression of Neurogenin-1 and −2 in human induced pluripotent stem cells (iPSC) triggers their high-efficiency differentiation into bipolar neurons (*17*) with TEs expression tightly regulated during the differentiation process (fig. S4A). We perturbed this system by either decreasing (via RNA interference) or increasing (via overexpression) the levels of the two KZFPs, or by repressing some of their HERVK and SVA targets with a CRISPR-based system (CRISPRi) (*18*). ZNF417/587-depleted iPSCs displayed a dysregulation of genes and TEs very reminiscent of that observed in hESCs (**Fig. 4A**). Neurons derived therefrom were characterized by the aberrant expression of non-neuronal genes (e.g. endothelium) (*19*) and the upregulation of transcripts related to potassium channel activity or to GABAergic neurotransmission (e.g. PRODH), whereas by contrast HERVK/SVA-CRISPRi-modified neurons displayed an induction of sodium and calcium channel-associated RNAs and a drop in GABAergic-related transcripts (**Fig. 4, B and C** and fig. S4, B and C). Furthermore, amongst 160 HERVKs predicted to encode for at least fragments of a retroviral envelope protein (ENV) recently demonstrated to be neurotoxic in the mouse brain and upregulated in cortical and anterior horn neurons of patients with sporadic amyotrophic lateral sclerosis (ALS) (*20*), we found fifteen to be targeted by ZNF417/587 and six of these to be upregulated upon *ZNF417/587* KD in iPSC-derived neurons (**Fig. 4D**). While ENV protein could not be easily detected in these cells, its induction was verified in NCCIT cells depleted for ZNF417/587 (**Fig. 4E**). We also observed an upregulation of the *np9* and *rec* alternative transcripts, which encode for proteins with oncogenic potential and promoting expression of the IFITM1 antiviral factor (*21*–*25*). As well, KZFP-depleted iN were characterized by the upregulation of IFNγ and ISGs such as TNF, IFITMs, APOBEC3B, IRF1, IFIH1 (a.k.a. MDA5), IFI44L, MOV10, RTP4, and Bst2 (**Fig. 4, F and G** and table S3) (*26*). This phenomenon was only partly abrogated by inhibiting the cytoplasmic DNA sensor STING (a.k.a.TMEM173) (fig. S4D), suggesting that it was not just due to ERV or SVA reverse transcripts but likely to additional TE-derived products as observed in *TREX1*- or *ADAR1*-inactivated astrocytes or neuronal progenitor cells, respectively, and upon *Rec* overexpression or treatment with inhibitors of DNA methyltransferases (*23, 27*–*31*). Reciprocally, levels of several ISG transcripts were decreased in HERVK-CRISPRi iN (**Fig. 4G**) or in ZNF417/587-overepressing iPSCs (fig. S4E). Finally, brain organoids derived from *ZNF417/587*-knockdown hESCs were smaller in size and displayed a greater abundance of PAX6-expressing immature cells than controls (**Fig. 4, H and I**), as well as increased levels of LTR/ERVK, SVAs and LTR/ERV1 RNA, alterations of neurotransmitter expression profiles and an inflammatory response reminiscent of that observed in KZFP-depleted pluripotent stem cells and neuron derivatives (**Fig. 4, J and K** and fig. S4F).

**Fig. 4:**
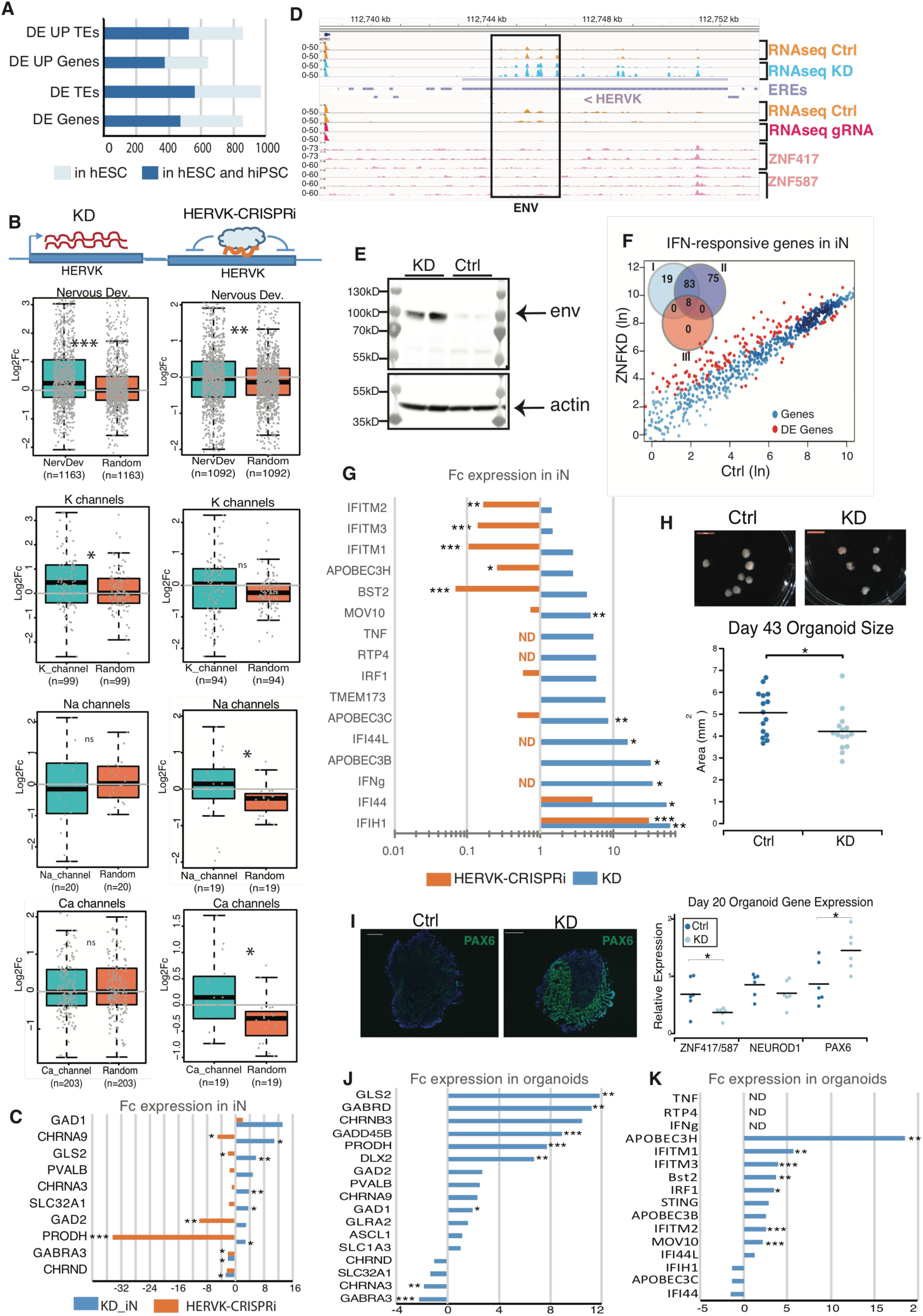
Impact of ZNF417/587 on neuronal differentiation, function and homeostasis. **(A)** Number of DE elements in *ZNF417/587* KD hESCs (light blue) and in common between hESCs+hiPSCs (dark blue). **(B)** Fold change in expression in KZFPs KD (left panels) or HERVK-silenced (right panels) *vs*. control induced neurons (iN) of genes related to each indicated category (as defined in Allen Brain Atlas GOs) compared to same number of random genes in each case. **(C)** Examples of genes DE in KZFPs KD and HERVK-CRISPRi iN and related to GABA-ergic pathway. **(D)** IGV screenshot of a region encompassing a full length HERVK encoding a consensus envelope with (top) RNAseq duplicates of control *vs*. KZFPs KD and control *vs*. HERVK-CRISPRi iN, and (bottom) ZNF417 and ZNF587 ChIPseq triplicates in H1 hESCs. **(E)** Western blot of independent duplicates of NCCIT-control or -KZFPs depleted cells lysates probed with HERVK anti-ENV antibody. Actin is used as loading control. **(F)** Dot-plot depicting expression of IFN-responsive genes (at least upregulated 5 times by IFN treatment in normal tissues or cells in Interferome database) in KZFPs KD *vs*. control iN. DE genes (Fc>2, FDR< 0.05) are highlighted in red. Venn diagram indicates number of DE genes stimulated by IFNs Type I, II or III. **(G)** Fold change expression of antiviral IFN-responsive genes in KZFPs KD and HERVK-CRISPRi *vs*. control iN (ND: not detected). **(H)** Brain organoids obtained 43 days post-differentiation of control or KZFPs KD hESCs, with size quantification below (Mann Whitney U-Test). **(I)** PAX6 immunostaining and quantification by RT-qPCR of indicated genes in organoids at 20 days post-induction of differentiation of indicated hESCs (two tailed T-test). **(J)** Fold change expression of neuronal function-related genes in KZFPs KD *vs*. control 20-days organoids. **(K)** Fold change expression of IFN-responsive genes in KZFPs KD *vs*. control 20-days organoids. For panels C, G, J, K, the average fold change of independent RNA-seq duplicates is shown (***FDR<0.001, **FDR<0.01, *FDR<0.05).

In sum, these results demonstrate that rather than just silencing TE-embedded regulatory sequences during early embryogenesis human KZFPs keep controlling their transcriptional impact later in development and in adult tissues. They further indicate that the evolutionary selection of some KZFP genes was key to the domestication of evolutionarily recent TEeRS towards the genesis of transcription networks active in the human brain. They finally imply that inter-individual differences in ZNF417, ZNF587 or their target TE-derived loci, many of which are species-specific and display some polymorphism in the human population, might translate into variations in brain development, function, and disease susceptibility.

## Acknowledgments

We thank Evarist Planet for the bioinformatics guidance, the EPFL genomics, Flow Cytometry and Histology Core Facilities for help with sequencing, immunohistochemistry and sorting, respectively. We thank Andrea Ablasser for the STING inhibitor, Raquel Fueyo and Joanna Wysocka for the NCCIT cell line and useful tips for the ENV detection, Riccardo Dainese for the help with the SMILE-seq experiment and the Tronolab for helpful discussions.

## Author contributions

T.P. and D.T. conceived the study, interpreted the data and wrote the manuscript. T.P. designed, performed and analyzed experiments. D.G. performed most of the bioinformatics analyses. C.R. performed most wet experiments; organoids-related experiments were performed by C.P. A.C., C.T. and J.D. developed bioinformatics tools. J.P. and P.E. conducted experiments. B.D. and V.B. gave intellectual input. All authors reviewed the manuscript;

## Data and materials availability

Sequencing data and processed files have been submitted to the NCBI Gene Expression Omnibus (GEO).

## Supplementary Materials

Materials and Methods

Figures S1-S4

Tables S1-S4

References (*31-47*)

