## Supplemental material for "Primate-restricted KRAB zinc finger proteins and target retrotransposons control gene expression in human neurons"

**This PDF file includes:**

Materials and Methods

Figs. S1 to S4

Tables S1 to S4

### Materials and Methods

Plasmids and Lentivectors production

Lentivectors with a 39bp sequence encompassing the HERVK14CI PBS were described previously[5]. The hairpin sequences against ZNF417/587 were designed from the Genetic perturbation portal (GPP) of the Broad institute and inserted with AgeI/EcoRI into pLKO.1.puro lentivector (SIGMA, TRCN0000018002). After assessing the specificity of various hairpins for depleting the 2 KZFPs, the shRNA targeting the sequence GCAGCATATTGGAGAGAAATT was used alone in all experiments except for the RNA sequencing in H1 hESCs where it was combined with one targeting AGTCGAAAGAGCAGCCTTATT in 2 of the 4 replicates. The pLKO.shLuc and pLKO empty vectors served as controls. Knock-down efficiency was verified by quantitative RT.PCR after puromycin selection and at the time of samples collection for each experiment. Lentivector containing Cas9KRAB and LTR5hs/SVAs sgRNA (ACCCCGTGCTCTCTGAAACAC ) was designed and validated as previously described[1] using a Cas9KRAB-expressing empty backbone as control. The ZNF417-HA and ZNF587-HA coding sequences from the previously described pTRE vectors[9] were swapped with the GFP cassette of the pRRL.PGK.GFP lentivector (http://www.addgene.org/, plasmid #12252) to generate KZFP-overexpressing vectors used in stem cells. Lentivectors production and titration protocols are detailed at http://tronolab.epfl.ch and lentiviral packaging plasmids are available at addgene (plasmids #12260, 12259). Viral titers were determined on HCT116 and equivalent number of transducing units were used for each vector in each experiment.

Cell culture

NCCIT cells were grown in RPMI-1640 (Thermo Fisher Scientific, Waltham, MA, USA), supplemented with 10% FBS, 1x Glutamax , 1x non-essential amino acids , and 1x antibiotic. Murine J1 ESCs, stably transduced with the diverse LV-HERVK vectors, were grown on gelatin, splitted with accutase, maintained in 2i + LIF media and transduced with the doxycycline-inducible pTRE-ZNFs lentivectors at a multiplicity of infection (MOI) of 60. Human iPSC and H1 hESC (WA01, WiCell) were maintained on hESC-qualified Matrigel (BD Biosciences) as previously described[5] and as approved by the Swiss Federal Office of Public Health and the Canton of Vaud Ethics committee. Stem cells were transduced at a MOI of 4, 15 or 100 for KD, CRISPRi or overexpression experiments respectively and selected with puromycin (0.5μg/ml) for 3 days for KD and CRISPRi. Puromycin-selected hiPSCs were differentiated in induced neurons by addition of doxycycline 1μg/ml for 4 days to induce expression of the exogenous Neurogenins as previously described[17]. Lyophilysed STINGi[32] was freshly reconstituted in DMSO, added at 0.5μM onto hiPSCs 45 minutes before transduction with KD lentivectors and replaced each day. Brain organoids were derived from puromycin selected control and KD H1 hESCs using the STEMdiff™ Cerebral Organoid Kit (#08570, StemCell Technologies) as per the manufacturers protocol, with day 0 denoting the transfer of hESCs to a 96 well plate for differentiation into embryoid bodies.

Proteins analyses

Proteins were extracted from 293T transfected with the pTRE plasmids with 150mM NaCl, 1%NP40 and 0,5% NaDoc and immunoprecipitated with either mouse IgGM (#I5381, SIGMA) or anti KAP1 (#MAB3662, millipore) antibodies. IP'ed material was loaded onto 4-20% SDS-PAGE and transferred to membranes for probing with anti-HA-HRPO (#12013819001, Roche) and rabbit anti-RBCC (#ab10483, abcam). Proteins were extracted from NCCIT with RIPA buffer, wettransferred overnight on PVDF membranes and blotted with rabbit anti-ENV (#HERM-1811-5, Austral Biologicals, 1/3000) and anti-Actin-HRPO (#ab20272, abcam) antibodies.

GFP reporter assay

Murine cells treated or not with doxycycline and transduced with pTRE-ZNFs or hES KD and control cells selected for puromycin resistance were plated in 12 well plates and transduced in duplicates with the same amount of transducing units for each LV-HERVK lentivector. GFP signal was read by flow cytometry.

RNA extraction, quantification and sequencing**.**

At least duplicates from 2 independent experiments were collected per sample. Total RNA was extracted either with TRIzol Reagent (Life Technologies), purified using the miRNeasy kit (Qiagen) and treated with RNase-Free DNase (Qiagen), or extracted with the high pure RNA isalation kit (Roche) with an on-column DNase treatment.

For RNA sequencing sample libraries were prepared using a TruSeq stranded mRNA sample preparation kit (Illumina). Libraries were sequenced on an Illumina Hi-Seq machine with stranded 75-base single or paired-end reads for the CRISPRi and ZNF KD RNAseqs respectively.

All RT-qPCR reactions were performed with RNAs from at least independent biological duplicates, using random hexamers and SuperScript II (Invitrogen) or Maxima H Minus, (thermo scientific) to generate cDNAs. Each cDNA was quantified in triplicates with SYBR green mix (Applied Biosystems). The -ΔΔCt method was used to calculate fold change. Negative controls without reverse transcription enzyme were processed in parallel. Primers used are described in Extended Data Table 3.

RNAseq mapping and analysis

For the Brainspan Atlas and GTEx RNAseq re-mapping, raw SRA files were downloaded from the dbGaP data portal using the prefetch tool from the NCBI SRA Toolkit v2.9.2 and then converted to FASTQ using the fastq-dump tool (with –split-3 for paired-end data).

Reads were mapped to the human genome (hg19) using HISAT2 (v2.1.0)[33] with default parameters. Genes counts were generated using uniquely mapped reads with HTSeq-count[34] (for KD hESC RNAseqs) or FeatureCounts[35] (iPS/iN and CRISPRi RNAseqs). TEs counts were generated using the uniquely mapped reads using the multiBamCov tool from the BEDtools software (for KD hESC RNAseqs) and featureCountss for all others. To avoid read assignation ambiguity between genes and TEs, a gtf file combining both features was provided to featureCounts. TEs reads in exons were always dismissed. Genes and TEs with low counts (at least as many reads as there are samples) were discarded. Normalization for sequencing depth was done for both genes and TEs using the TMM method as implemented in the limma package of Bioconductor[36] and using the counts on genes as library size. Differential gene expression analysis was performed using voom[37] as it has been implemented in the limma package of Bioconductor. A moderated t-test (as implemented in the limma package of R) was used to test significance. *P* values were adjusted for multiple testing using the Benjamini-Hochberg's method. A gene (or TE) was considered to be differentially expressed when the absolute fold change between groups was bigger than 2 and the FDR (adjusted *P* value) was smaller than 0.05 (***$P<0.001$, **$P<0.01$, and *$P<0.05$). Interspecies RNA-seq normalization was performed as previously described[1].

Repeats merging and aging

The RepeatMasker 4.0.5 (Library 20140131, http://www.repeatmasker.org/species/hg.html) was used to generate an in-house repeats list where fragmented annotated ERVs have been reassembled in one single element when relevant and according to the following procedure: ERVs fragments are annotated either as fragmented internal parts (ERV-int) or LTRs (Long Terminal Repeats). Starting from each annotated ERV-int element, we first computed a frequency table of the surrounding retroelements fragments. We then computed the frequencies distributions of each LTR type for each ERV–int sub-family, and performed a Wald test to assign a p-value to each LTR/ERV-int pair. To merge a LTR to an ERV-int fragment, it had to represent more than 2% of the pairs, the p-value had to be smaller than 0.001, the two elements had to be in the same orientation and the distance between the two elements had to be shorter than 100bp. The name of the internal part was given to the resulting LTR/ERV-int merged element. For example, an element resulting from LTR7 and HERVH-int fragments merging is identified as HERVH-int. Relevance of LTRs and internal pieces merging was verified when possible in the literature and in the DFAM database of repetitive DNA based on profile hidden Markov models (dfam.org). Fragmented ERV-int or fragmented LTRs from the same sub-family (same name in Repbase database), with the same orientation and closer than 100bp were also merged.

We used RNA-sequencings with a depth superior to 30-40 million reads and at least 75 bp in length as we determined, by comparing various RNA-seq approaches for their respective yield in mappable TE transcripts, that it allowed us to assign more than 91% of repeats-derived reads to specific unique genomic loci through our TE-optimized analytical pipeline.

TE age was estimated by measuring substitution density of TE loci by comparison with their respective subfamily consensus. Substitutions were then separated between transitions and transversions to feed a Kimura 2-parameters model[38] (in-house script). To estimate an evolutionary age from Kimura substitution levels, we assumed a constant rate of substitution per million years of 2.2e-3 as described previously[39] which allowed to get divergence time in million years for each TE loci.

ChIP-seq

KAP1 ChIPseq has been described earlier[5]. Chromatin for ZNF-HA ChIPs was prepared in triplicates from $3.10^7$ cells and chromatin for histones ChIPs in at least duplicates from 2 independent experiments with $10^7$ cells, as followed: adherent stem cells were detached with TrypLE, diluted in DMEM KO medium and washed with PBS. Cells were fixed with methanol-free formaldehyde (333 mM final) 10min at room temperature, diluted with Tris pH8 (200 mM final), washed 2 times with cold PBS and pellets were frozen at -80°C. Cross-linked cells were incubated 10min at 4°C in buffer I (50 mM HEPES-KOH pH 7.4, 140 mM NaCl, 1 mM EDTA, 0.5 mM EGTA, 10% Glycerol, 0.5% NP40, 0.25% Tx100, supplemented with protease inhibitors), spinned, incubated 10min at 4°C in buffer II (10 mM Tris pH8, 200 mM NaCl, 1 mM EDTA, 0.5 mM EGTA, proteases inhibitors), spinned, washed 3 times at 4°C in buffer III (buffer II supplemented with 0.1% NaDOC and 0.25% NLS) and resuspended in 1ml buffer III before being transferred to a TC12X12 AFA fiber tube (Covaris). Sonication was performed with an automated Covaris Focus ultra-sonicator with the following parameters: 20min, 5% duty cycle, 140W, 200 cycles and extracts were cleared by centrifugation 10min at 4°C. Chromatin sonication was evaluated on a Bioanalyser DNA chip (Agilent) with one aliquot of material treated with RNAse A O/N at 65°C and purified with Qiagen minElute columns. Immunoprecipitations were performed in low DNA binding tubes using Dynabeads® (ThermoFisher) after a pre-clearing step for histones ChIPs and as recommended by the manufacturer. Antibodies specific for HA

(#901503, BioLegend), H3K9me3 (#C15410056, Diagenode), H3K4me1 (#pAb-037-050, Diagenode) and H3K27ac (#ab4729, abcam) were used. IPed material was eluted and treated with proteinase K ON at 65°C in EB (TE 1X, 1% SDS), and purified on Qiagen minElute columns. Total inputs and ChIPed samples were prepared for libraries using 10ng of material. Aliquots of the samples were tested in qPCR using the KAPA SYBR FAST qPCR Master Mix (KAPA Biosystems, KK4604) to determine the optimal number of PCR cycles needed to amplify each library without reaching saturation. Libraries were amplified with KAPA HiFi Hotstart Readymix polymerase (KAPA Biosystems, KK2602), size-selected using Ampure XP beads (Beckman Coulter) and always quality checked on Bioanalyser DNA high sensitivity chip (Agilent) and quantified by Qubit ds DNA HS assay kit (Qubit® 2.0 Fluorometer, Invitrogen). ChIP enrichment was always confirmed by qPCR with positive versus negative controls, before and after libraries preparation. Libraries were sequenced with 75bp single or paired-end reads on an Illumina Hi-Seq machine with standard Illumina protocols.

ChIP-seq analysis

The previously published KAP1 ChIPseq[5] has been remapped with our new pipeline. Reads were mapped to the human genome (hg19) using the short read aligner program Bowtie2, with the sensitive local mode (the exact parameters are: bowtie2 -p 6 -t --sensitive-local -x $index -U $reads). Only reads with a mapping quality score higher than 10 were retained.

Before peak calling, bam files were filtered removing reads aligning to ENCODE blacklist regions and to regions of ChIP experiments with high signal in the input using Bioconductor package GreyListChIP (https://bioconductor.org/packages/GreyListChIP). Peaks for histone marks were called via EPIC (reimplementation of SICER[40]) keeping only peaks with an FDR below 1%. Peaks for ZNF417, ZNF587 and KAP1 were called using MACS2[41] (with - -BAMPE option as from paired end reads), keeping only peaks with score higher than 80. Peaks from biological replicates were merged using the Bioconductor package DiffBind (https://bioconductor.org/packages/DiffBind) and when triplicates were made, only peaks that overlapped in at least 2 peak sets were kept and merged into consensus peaks.

The intersectBed tool (with default parameters and unique option) from the BEDTools suite[42] was used to calculate intersections, and the genomeCoverageBed tool to generate coverage files, which were converted into bigWig files with the bedGraphToBigWig tool (provided by UCSC).

Multiple sequence alignment plot

Fasta sequences from KZFP overlapping ERVK integrants were extracted from the hg19 genome assembly and processed as described previously[43] except that regions in the alignment consisting of more than 90% of gaps were trimmed out. For each aligned integrant, the ZNF417, ZNF587, KAP1 or H3K9me3 ChIP-seq signal was extracted from the BAM alignment files using the python pysam library and scaled to the interval [0,1] before being superimposed to the alignment. The average ChIP-seq signals were plotted on top.

SMILE-seq with methylated probes

SMILE-seq device and analysis was previously described[12]. Motif enrichment analysis was performed using HOMER[44] . Methylated and unmethylated probes were used here.

Human genetic data

Information on human genetic data and LoF muations was obtained from The Genome Aggregation Database (gnomAD)[10,45] (release-2.0.2), containing exome and whole genome sequencing data for 123,136 and 15,496 individuals, respectively.

Immunofluorescence on brain organoids

Day 20 organoids were fixed with 4% PFA pH 7.4 for 15 minutes, washed 2 times with PBS and placed in 30% sucrose solution overnight at 4°C. Fixed organoids were placed in cryomolds, covered with cryomatrix embedding resin (#6769006, Thermo Scientific), snap frozen with isopentae, followed by cutting 8μm sections with a cryostat (Leica) to Superfrost Plus slides (Thermo Scientific). For immunoflurorescent staining, tissue was permeablized by 0.25% Triton X-100 for 10 minutes, washed 3x with PBS, blocked for 30 minutes in 1% BSA in PBS, incubated with primary antibody (PAX6 **#561462,** BD Biosciences) overnight at 4°C then incubated with secondary Alexafluor antibody (Thermo Scientific) for 40 minutes. Nuclear staining was performed with DAPI and slides were mounted with FluoromountG (#00-4958-02, Thermo Scientific). Fluorescent imaging was performed with a Leica DM5500 microscope. For widefield organoid images, a Leica stereoscope was used and the area of individual organoids were measured with ImageJ.

Allen Brain transcriptome mapping

Only uniquely mapped reads and TEs with more than 50 reads were retained for analysis. The package used to generate the plots was previously described[46].

Statistical analysis

R version 1.1.447 and Wilcoxon test were used for statistical analyses of boxplots. The details of the other statistical tests have been described in the figure legends or in the respective methods sub-sections. In all analyses, ***$P$<0.001, **$P$<0.01, and *$P$<0.05.

A

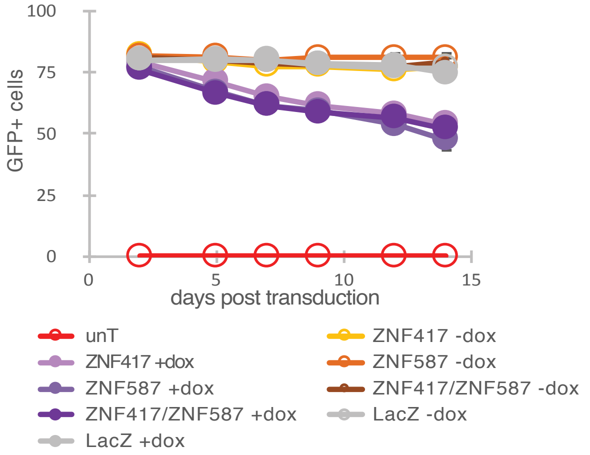


B

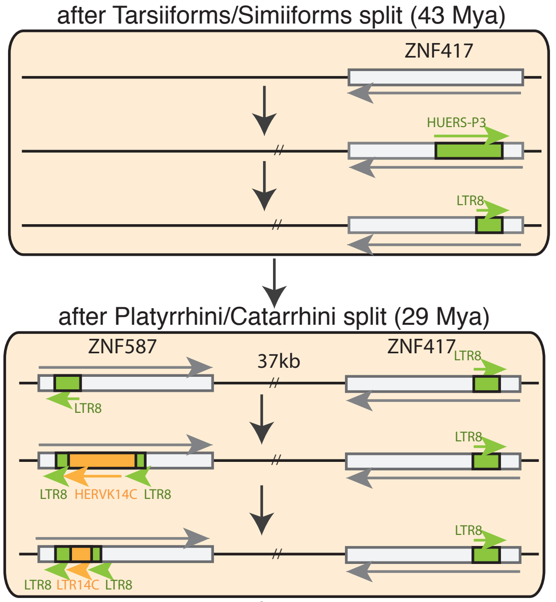


human genome

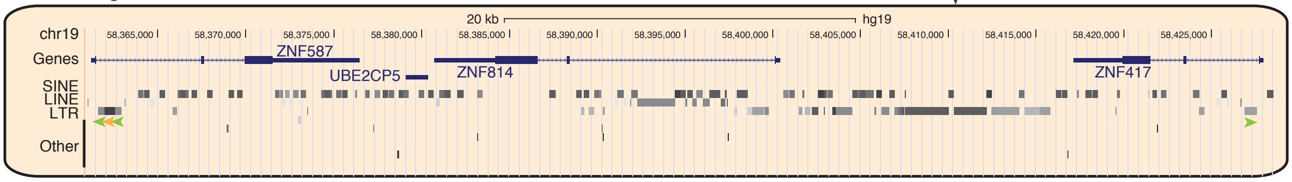


C

| | species | age (My) | ZF1 | ZF2 | ZF3 | ZF4 | ZF5 | ZF6 | ZF7 | ZF8 | ZF9 | ZF10 | ZF11 | ZF12 | ZF13 | ZF14 | ZF15 | |
|---|---|---|---|---|---|---|---|---|---|---|---|---|---|---|---|---|---|---|
| ZNF587 | Homo sapiens | 0 | DHD | XXX | RSN | RSQ | QSS | QNQ | QCN | QNH | YHE | RHV | NSI | SAV | ESK | RTH | XXX | hominoids |
| ZNF417 | Homo sapiens | 0 | DHD | XXX | RSN | RSQ | QSS | XXX | QCN | QNQ | YHE | RHV | NCI | SAV | ESK | RTH | XXX | |
| ZNF587 | Pan troglodytes | 6.6 | DHD | XXX | RSN | RSQ | QSS | XXX | QCN | QNH | YHE | RHV | NSI | SAV | ESK | RTH | XXX | |
| ZNF417 | Pan troglodytes | 6.6 | DHD | - | RSN | RSQ | QSS | XXX | QCN | QNQ | YHE | RHV | NCI | SAV | - | RTH | (XXX) | |
| ZNF587 | Pongo abelii | 15.8 | (DHD) | - | (XXX) | (XXX) | (QSS) | (QNQ) | (QCN) | (QNH) | (YHE) | - | - | (SAV) | (ESK) | (RTH) | (XXX) | |
| ZNF417 | Pongo abelii | 15.8 | DHD | XXX | RSN | RSQ | QSS | XXX | QCN | QNQ | YHE | RHV | NCI | SAV | ESK | RTH | XXX | |
| ZNF587 | Nomascus leucogenys | 19.9 | DHD | - | RSN | QSR | QSS | QNQ | QCN | QNH | YHE | RHV | NSI | SAV | ESK | RTH | XXX | Old world monkeys |
| ZNF417 | Nomascus leucogenys | 19.9 | DHD | - | RSN | RSQ | QSS | QNQ | QCN | QNQ | YHE | RHV | NCI | SAV | ESK | RTH | XXX | |
| ZNF587? | Chlorocebus sabaeus | 29.1 | - | - | - | - | - | - | (QCN) | (QCN) | (XXX) | (RHV) | (SSI) | (SAA) | (ESK) | (RTH) | (ESK) | |
| ZNF417 | Chlorocebus sabaeus | 29.1 | - | - | - | - | - | (QNQ) | (QCN) | (QNQ) | (YHE) | (RHV) | (NSI) | (SAV) | (ESK) | (RTH) | (ESK) | |
| ZNF417 | Papio anubis (olive baboon) | 29.1 | DHE | XXX | RSN | RSQ | XXX | QKQ | QCN | QNQ | YHE | RHV | NSI | SAV | ESK | RTH | ESK | |
| ZNF417 | Papio anubis (olive baboon) | 29.1 | DHE | XXX | RSN | RSQ | XXX | QKQ | QCN | QNQ | YHE | RHV | NSI | SAV | ESK | RTH | XXX | |
| ZNF417 | Rhinopithecus roxellana | 29.1 | DHD | XXX | RSK | QSQ | XXX | QKQ | - | QNQ | YHE | RHV | NSI | (XXX) | ESK | - | (ESK) | |
| ZNF417 | Rhinopithecus roxellana | 29.1 | HHD | - | RSK | - | QCN | QNQ | QCN | QNQ | YHE | RHV | SSI | SAV | ESK | RTH | ESK | |
| ZNF417 | Colobus angolensis palliatus | 29.1 | DHD | XXX | RSK | QSL | XXX | QKQ | QCN | QNQ | YHE | RHV | NSI | SAV | ESK | RTH | XXX | |
| ZNF417 | Papio hamadryas | 29.1 | (DHE) | (XXX) | (RSN) | (RSQ) | (XXX) | (QKQ) | (QCN) | (QNQ) | (YHE) | (RHV) | (NSI) | (SAV) | (ESK) | (RTH) | (ESK) | |
| ZNF417 | Macaca fascicularis | 29.1 | DHE | XXX | RSN | QSQ | QRN | QKQ | QCN | QNQ | YHE | RHV | NSI | SAV | ESK | RTH | (ESK) | |
| ZNF417 | Macaca mulatta lasiota | 29.1 | (DHE) | (XXX) | (RSN) | (QSQ) | (QRN) | (QKQ) | (QCN) | (QNQ) | (YHE) | (RHV) | (NSI) | (SAV) | (ESK) | (RTH) | (ESK) | |
| ZNF417 | Macaca nemestrina | 29.1 | DHE | XXX | RSN | QSQ | QRN | QKQ | QCN | QNQ | YHE | RHV | NSI | SAV | ESK | RTH | ESK | |
| ZNF417 | Cercocebus atys | 29.1 | DHE | XXX | RSN | QSQ | QRN | QKQ | QCN | QNQ | YHE | RHV | NSI | SAV | ESK | RTH | ESK | |
| ZNF417 | Mandrillus leucophaeus | 29.1 | DHE | XXX | RSN | QSQ | QRN | QKQ | QCN | QNQ | YHE | RHV | NSI | SAV | ESK | RTH | ESK | |
| ZNF417 | Nasalis larvatus | 29.1 | (XXX) | - | (RSK) | (QSQ) | (XXX) | (QKQ) | (QCN) | (QNQ) | (YHE) | (RHV) | (NSI) | (SAV) | (ESK) | (RTH) | (ESK) | |
| ZNF417 | Nancy Ma's night monkey | 43.1 | - | XXX | KSN | QHQ | RNL | QNL | - | QNQ | YHE | KHV | NCL | YAV | ESK | RAQ | - | NWM |

D

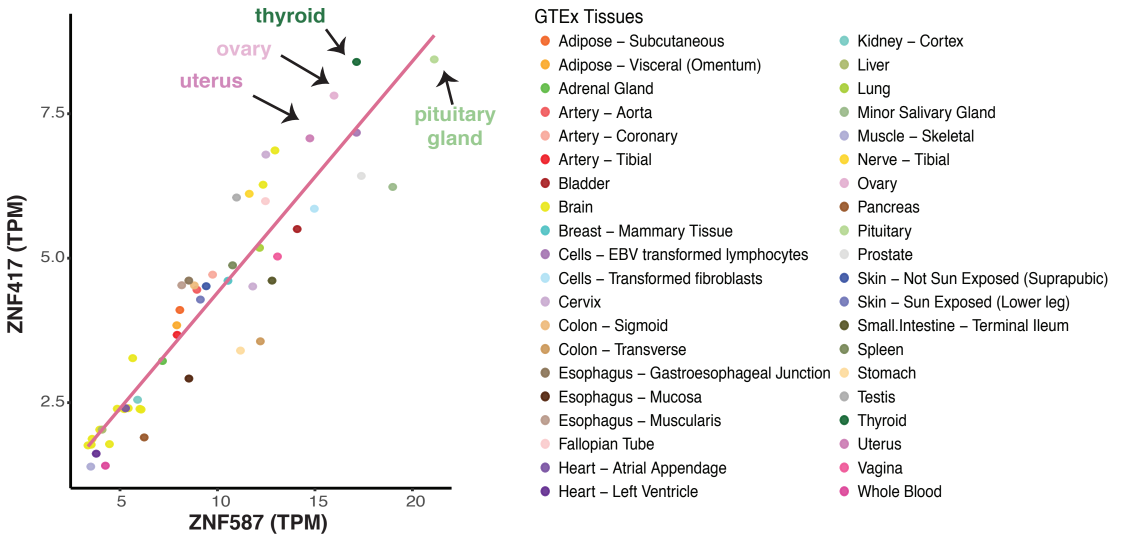

**Fig. S1. The 2 TE controllers arose from duplication in Primates and are co-expressed in many tissues**

**(A)** The doxycycline induction of either ZNF417, ZNF587 (or both together) in mESCs is triggering the silencing of the HERVK-R containing LV. Cells expressing an inducible LacZ protein were used as controls. Average and s.d. of duplicates are depicted. UnT: untransduced. **(B)** Skim illustrating evolution of the *ZNF417/ZNF587* genomic locus, with appearance of the ancestral gene in Simiiforms and its duplication in Catarrhini followed by the colonization of the ZNF587 by the HERVK14C, and a UCSC screenshot of the human genomic locus. **(C)** Evolution of ZNF417- and ZNF587-orthologs zinc fingerprints from the ancestral to the human genes. Fingerprint elements similar or different between the 2 human proteins are highlighted in red and green respectively; XXX indicate residual zinc fingers with mutated Cys or His in C2H2 structural motif. Parentheses indicate sequences not found in current annotations. **(D)** ZNF417 and ZNF587 expression across 40 distinct tissues based on GTEx data (v7). TPM: transcripts per million.

Fig. S2

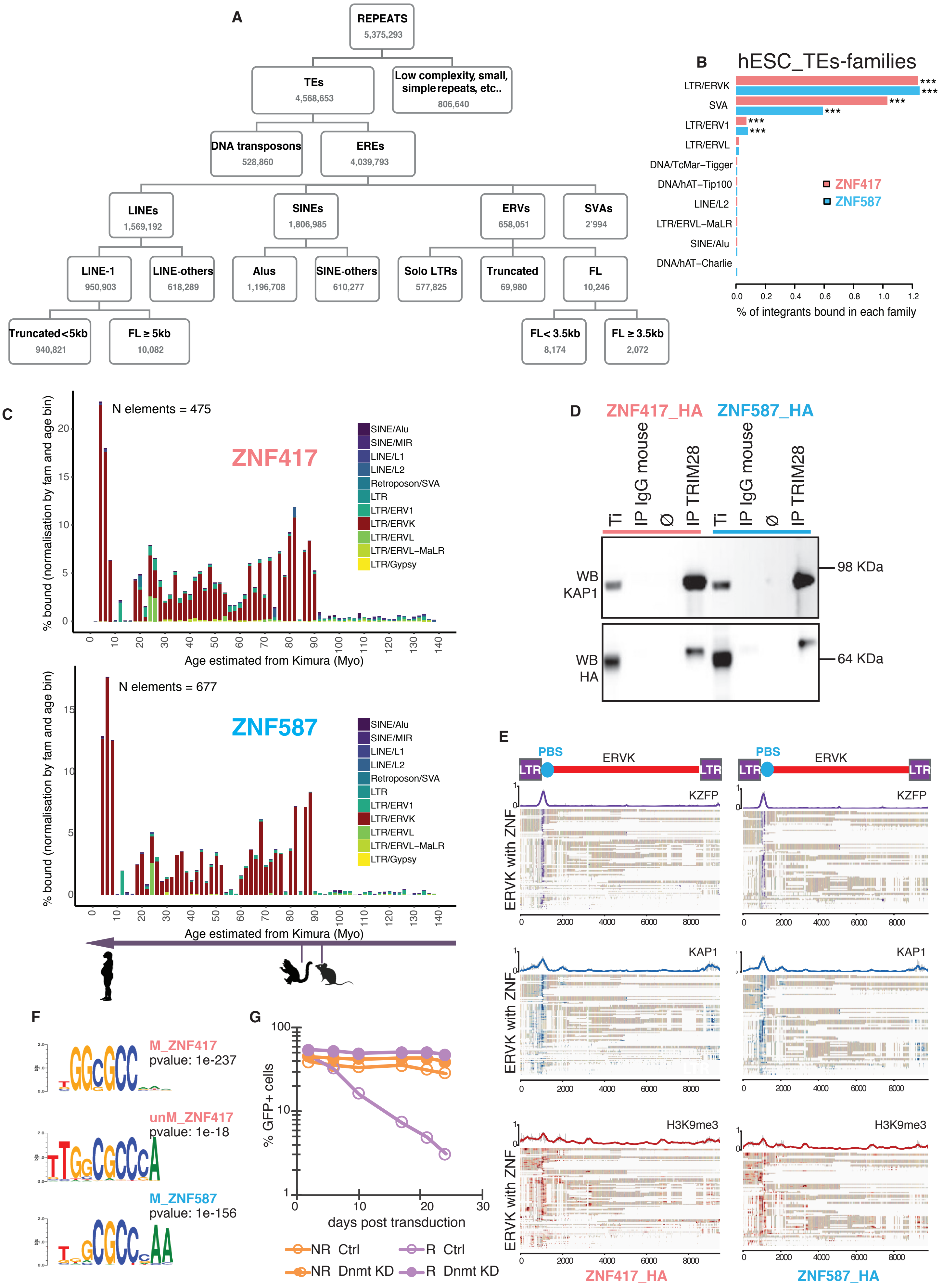
A
REPEATS 5,375,293
TEs 4,568,653
Low complexity, small, simple repeats, etc.. 806,640
DNA transposons 528,860
EREs 4,039,793
LINEs 1,569,192
SINEs 1,806,985
ERVs 658,051
SVAs 2'994
LINE-1 950,903
LINE-others 618,289
Alus 1,196,708
SINE-others 610,277
Solo LTRs 577,825
Truncated 69,980
FL 10,246
Truncated<5kb 940,821
FL ≥ 5kb 10,082
FL< 3.5kb 8,174
FL ≥ 3.5kb 2,072
B
hESC_TEs-families
LTR/ERVK
SVA
LTR/ERV1
LTR/ERVL
DNA/TcMar–Tigger
DNA/hAT–Tip100
LINE/L2
LTR/ERVL–MaLR
SINE/Alu
DNA/hAT–Charlie
ZNF417
ZNF587
% of integrants bound in each family
C
N elements = 475
ZNF417
N elements = 677
ZNF587
% bound (normalisation by fam and age bin)
Age estimated from Kimura (Myo)
SINE/Alu
SINE/MIR
LINE/L1
LINE/L2
Retroposon/SVA
LTR
LTR/ERV1
LTR/ERVK
LTR/ERVL
LTR/ERVL–MaLR
LTR/Gypsy
D
ZNF417_HA
ZNF587_HA
Ti
IP IgG mouse
Ø
IP TRIM28
WB KAP1
WB HA
98 KDa
64 KDa
E
PBS
ERVK
LTR
KZFP
KAP1
H3K9me3
ERVK with ZNF
ZNF417_HA
ZNF587_HA
F
M_ZNF417
pvalue: 1e-237
unM_ZNF417
pvalue: 1e-18
M_ZNF587
pvalue: 1e-156
G
% GFP+ cells
days post transduction
NR Ctrl
R Ctrl
NR Dnmt KD
R Dnmt KD

**Fig. S2. Primate-specific TEs are epigenetically silenced by the KAP1 recruiting KZFPs**

**(A)** Summary of our improved version of Repbase (db20140131, see Methods). Full length (FL) ERVs are elements with internal sequences flanked by 2 LTRs. FL LINE-1 are elements ≥5kb in length. Only SVA sequences longer than 200bp are considered as *bona fide* integrants. **(B)** Percentage of integrants from indicated TE families overlapping with ZNF417 and ZNF587 peaks as obtained by ChIP-seq triplicates in hES H1 cells. Only families with at least 5 integrants bound are shown. (****P*<0.001, hypergeometric test). **(C)** Percentage of bound elements per family and per age as determined by Kimura score. **(D)** KAP1 associates with HA-tagged versions of ZNF417 and ZNF587 as revealed by KAP1 immunoprecipitation followed by western blotting using anti-HA or -KAP1 antibodies. **(E)** Multiple sequence alignement plot of KZFPs (top), KAP1 (middle) and H3K9me3 (bottom) ChIP-seq signal enrichment in hESCs on LTR/ERVK loci bound by indicated KZFP, marking the PBS-coding region at the 5' end of the integrants. Each row is independently normalized with darkness of color proportionate to enrichment. LTR/ERVK sequences are aligned, and white boxes correspond to gaps in the alignments. **(F)** ZNF417 and ZNF587 binding motifs obtained with unmethylated (unMe) or methylated (M) SMILE-seq probes. We did not see any specific motif enrichment for ZNF587 with unmethylated sequences. **(G)** Silencing assay of the depicted HERVK-containing PGK-GFP LVs in either control (Ctrl) or Dnmt3A/B depleted hESCs (Dnmt KD). UnT: untransduced negative control. Data are representative of two independent biological replicates.

Fig. S3

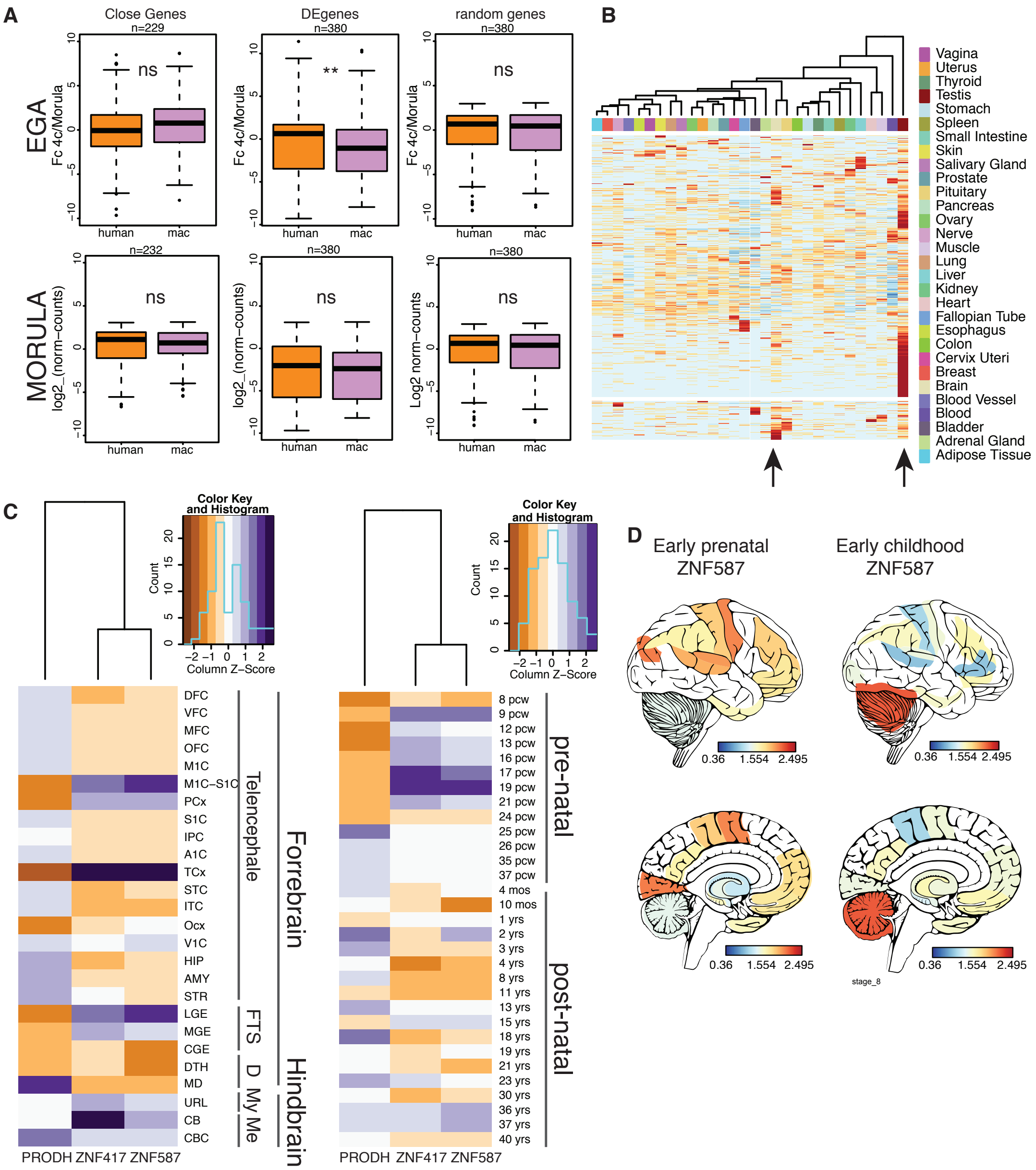
A
Close Genes
n=229
DEgenes
n=380
random genes
n=380
EGA
Fc 4c/Morula
ns
**
ns
human
mac
MORULA
log2_(norm−counts)
Log2 norm−counts
n=232
n=380
n=380
B
Vagina
Uterus
Thyroid
Testis
Stomach
Spleen
Small Intestine
Skin
Salivary Gland
Prostate
Pituitary
Pancreas
Ovary
Nerve
Muscle
Lung
Liver
Kidney
Heart
Fallopian Tube
Esophagus
Colon
Cervix Uteri
Breast
Brain
Blood Vessel
Blood
Bladder
Adrenal Gland
Adipose Tissue
C
Color Key and Histogram
Count
Column Z−Score
DFC
VFC
MFC
OFC
M1C
M1C−S1C
PCx
S1C
IPC
A1C
TCx
STC
ITC
Ocx
V1C
HIP
AMY
STR
LGE
MGE
CGE
DTH
MD
URL
CB
CBC
Telencephale
FTS
D
My
Me
Forrebrain
Hindbrain
PRODH ZNF417 ZNF587
8 pcw
9 pcw
12 pcw
13 pcw
16 pcw
17 pcw
19 pcw
21 pcw
24 pcw
25 pcw
26 pcw
35 pcw
37 pcw
4 mos
10 mos
1 yrs
2 yrs
3 yrs
4 yrs
8 yrs
11 yrs
13 yrs
15 yrs
18 yrs
19 yrs
21 yrs
23 yrs
30 yrs
36 yrs
37 yrs
40 yrs
pre-natal
post-natal
D
Early prenatal ZNF587
Early childhood ZNF587
0.36 1.554 2.495
stage_8

**Fig. S3. ZNF417/587 targeted TEs–enhancers control genes in early embryos and later in development**

**(A)** Expression of close-, DE- or random- human and macaque gene orthologs during early genome activation stages (EGA, 4 cell to morula, upper panels) or at the morula stage (lower panels). **(B)** HeatMap of average expression of ZNF417- and ZNF587- bound TEs across tissues using GTEX data and unsupervised clustering. Arrows indicate brain and testis. **(C)** Average developmental stages expression in the various substructures of the brain (left panel) or average substructures expression in the various developmental stages (right panel) of *ZNF417, ZNF587* and *PRODH* based on BrainSpan Atlas of the Developing Human Brain. **(D)** Spatial representation of *ZNF587* expression.

**Fig. S4**

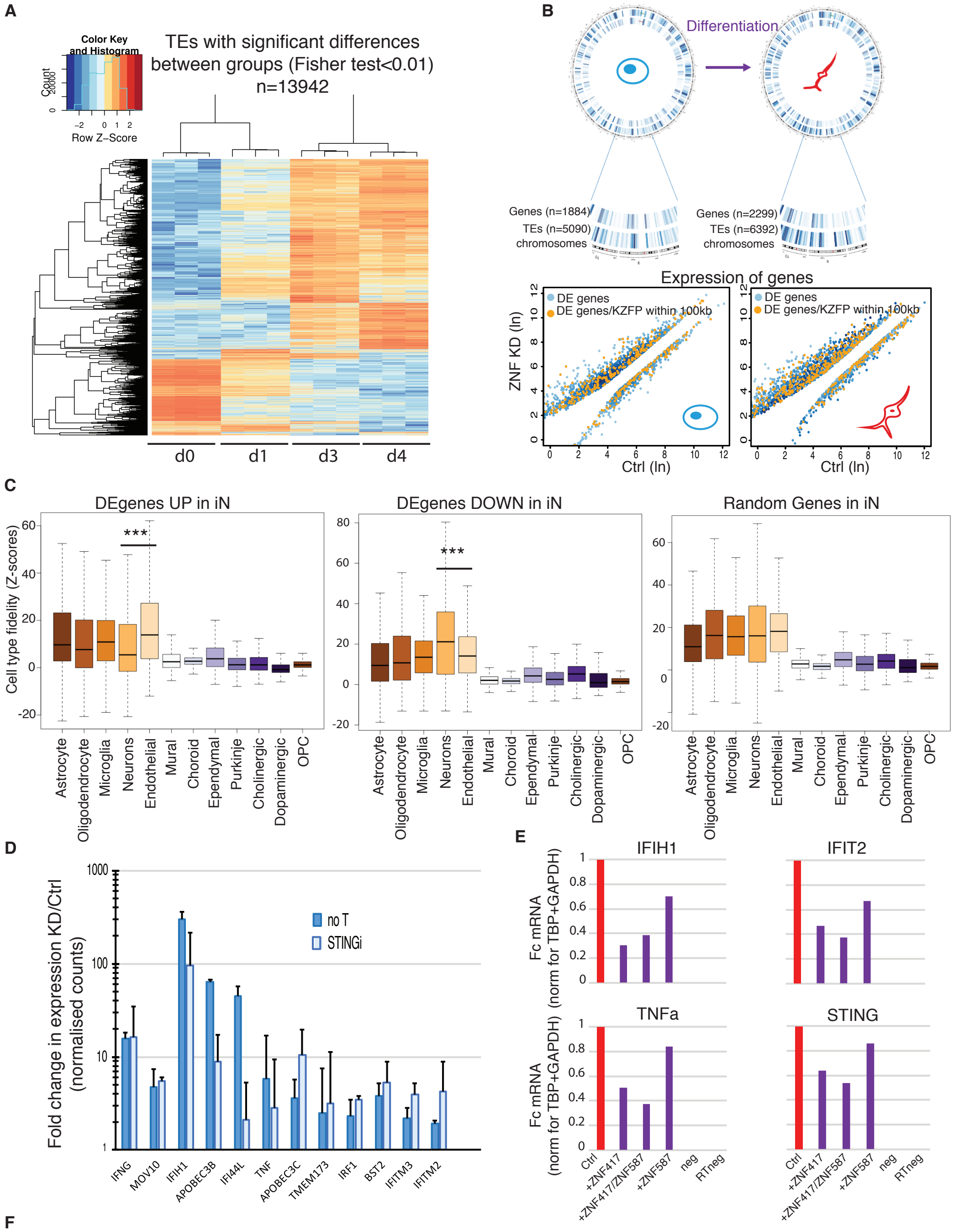


**F**

| class.fam | Expressed | DiffExpress.count | DiffExpress.Prop | DiffExpress.pval | DownRegul.count | DownRegul.Prop | DownRegul.pval | UpRegul.count | UpRegul.Prop | UpRegul.pval |
|---|---|---|---|---|---|---|---|---|---|---|
| LTR/ERV1 | 799 | 161 | 20.2 | 1.13E-12 | 77 | 9.6 | 0.008747067 | 84 | 10.5 | 4.93E-13 |
| Retroposon/SVA | 95 | 31 | 32.6 | 5.25E-08 | 5 | 5.3 | 0.837690796 | 26 | 27.4 | 4.93E-13 |
| LTR/ERVK | 188 | 36 | 19.1 | 0.001874709 | 13 | 6.9 | 0.634097316 | 23 | 12.2 | 9.41E-05 |
| LTR/ERVL-MaLR | 760 | 136 | 17.9 | 1.77E-07 | 85 | 11.2 | 6.48E-05 | 51 | 6.7 | 0.012775211 |
| LTR/ERVL | 404 | 54 | 13.4 | 0.161749645 | 32 | 7.9 | 0.35899835 | 22 | 5.4 | 0.949548253 |
| SINE/Alu | 5023 | 348 | 6.9 | 1 | 211 | 4.2 | 1 | 137 | 2.7 | 1 |
| LINE/L1 | 3270 | 477 | 14.6 | 9.73E-09 | 351 | 10.7 | 1.69E-15 | 126 | 3.9 | 1 |
| SINE/MIR | 1877 | 220 | 11.7 | 0.492314024 | 139 | 7.4 | 0.48486194 | 81 | 4.3 | 1 |
| LINE/L2 | 1616 | 173 | 10.7 | 0.910011627 | 102 | 6.3 | 0.962565226 | 71 | 4.4 | 1 |

**Fig. S4. The KZFPs-mediated control of TEs is necessary for induced neurons and brain organoids *in vitro* differentiation**

**(A)** HeatMap depicting expression of TEs with significant differences between days 0, 1, 3 and 4 of hiPSCs to iN differentiation. **(B)** Upper panel: DE genes and TEs in control versus KD hiPSCs and iN are placed along human chromosomes. The zoom shows the degree of overlap between differentially expressed TEs and genes. Lower panel: MA-plot of DE genes in control versus KD iPSCs and differentiated iNs. DE genes located within 100kb of a ZNF417 or ZNF587 peak are highlighted in orange. **(C)** Boxplots of cell type fidelity Z-scores, as described in Kelley et al19, of genes up- (left panel) or down-regulated (middle panel) in KD cells and random genes (right panel). **(D)** Average fold change in expression, as determined by RNAseq with independent duplicates of IFN-stimulated genes in control vs. KZFPs depleted iN, in non treated and STINGi-treated cells. **(E)** Fold change in expression, as determined by RT-qPCR with independent duplicates, of IFN-responsive genes in control cells vs. hESCs overexpressing ZNF417, ZNF587 or both. RTneg: DNA contamination control reaction, performed without reverse transcriptase. Neg: negative control with no RNA input. **(F)** Expression of indicated TE family members in control versus KZFPs-depleted organoids. Counts, proportion and *P* values of expressed, DE, downregulated and upregulated elements are indicated.

**Table S1.**

Human specific LTR/ERVK bound by ZNF417 or ZNF587.

**Human specific LTR/ERVK bound by ZNF417 or ZNF587**

| nb | chr | start | end | merged TEs | size | strand | Fam | subfam merged name | length | substitution_level | Kimura age | Consensus ENV | polymorphic[11] |
|---|---|---|---|---|---|---|---|---|---|---|---|---|---|
| 1 | chr3 | 112743124 | 112752282 | LTR5_Hs\|HERVK-int\|HERVK-int\|LTR5_Hs | 9159 | - | LTR/ERVK | HERVK-int | 9158 | 0.33 | 1.5 | Yes | Yes |
| 2 | chr3 | 148281477 | 148285396 | HERVK-int | 3920 | - | LTR/ERVK | HERVK-int | 3919 | 0.45 | 2.04545455 | No | nd |
| 3 | chr6 | 78426662 | 78436083 | LTR5_Hs\|HERVK-int\|LTR5_Hs | 9422 | - | LTR/ERVK | HERVK-int | 9421 | 0.46 | 2.09090909 | Yes | Yes |
| 4 | chr1 | 156149014 | 156149981 | LTR5_Hs | 968 | + | LTR/ERVK | LTR5_Hs | 967 | 0.55 | 2.5 | No | nd |
| 5 | chr22 | 18926187 | 18935361 | LTR5_Hs\|HERVK-int\|HERVK-int\|LTR5_Hs | 9175 | + | LTR/ERVK | HERVK-int | 9174 | 0.690764778 | 3.1398399 | yes | nd |
| 6 | chr6 | 31952469 | 31958829 | LTR14\|HERVKC4-int\|LTR14 | 6361 | - | LTR/ERVK | HERVKC4-int | 6360 | 0.715539478 | 3.25245217 | No | CNV |
| 6 bis | chr6 | 31985207 | 31991567 | LTR14\|HERVKC4-int\|LTR14 | 6361 | - | LTR/ERVK | HERVKC4-int | 6360 | 0.715539478 | 3.25245217 | No | CNV |
| 7 | chr10 | 27182399 | 27183366 | LTR5_Hs | 968 | + | LTR/ERVK | LTR5_Hs | 967 | 0.75 | 3.40909091 | No | Yes |
| 8 | chr7 | 4622057 | 4640031 | LTR5_Hs\|HERVK-int\|LTR5_Hs\|HERVK-int\|LTR5_Hs | 17975 | - | LTR/ERVK | HERVK-int | 17974 | 0.76 | 3.45454546 | yes | Yes |
| 9 | chr3 | 185280336 | 185289515 | LTR5_Hs\|HERVK-int\|HERVK-int\|LTR5_Hs | 9180 | - | LTR/ERVK | HERVK-int | 9179 | 0.96 | 4.36363636 | Yes | nd |
| 10 | chr12 | 111007843 | 111009325 | LTR5_Hs\|HERVK-int | 1483 | + | LTR/ERVK | HERVK-int | 1482 | 1.045705604 | 4.75320729 | No | nd |
| 11 | chr7 | 104389349 | 104393266 | HERVK-int | 3918 | - | LTR/ERVK | HERVK-int | 3917 | 1.06 | 4.81818182 | No | nd |
| 12 | chr7 | 30757734 | 30757871 | LTR5B | 138 | + | LTR/ERVK | LTR5B | 137 | 1.47 | 6.68181818 | No | nd |

nd: not described as polymorphic

**Table S2.**
Normalized counts for DE genes (Fc<2 or Fc>2, FDR>0.05) in hES H1 KD cells

**Table S3.**
Normalized counts for IFN-responsive (interferome database) coding genes in iN.

**Table S4.**
Primers used in this study.
